# Network size affects the complexity of activity in human iPSC-derived neuronal populations

**DOI:** 10.1101/2023.10.31.564939

**Authors:** Yavuz Selim Uzun, Renata Santos, Maria C. Marchetto, Krishnan Padmanabhan

**Affiliations:** Department of Physics and Astronomy, University of Rochester; Del Monte Institute for Neuroscience, University of Rochester School of Medicine; Université Paris Cité, Institute of Psychiatry and Neuroscience of Paris (IPNP), INSERM U1266, Signaling mechanisms in neurological disorders, 102 rue de la Santé, 75014 Paris, France; Institut Imagine, INSERM U1163, Mechanisms and therapy of genetic brain diseases, Université Paris Cité, 24 Boulevard du Montparnasse, 75015 Paris, France; Institut des Sciences Biologiques, CNRS, 16 rue Pierre et Marie Curie, 75005 Paris, France; Department of Anthropology, University of California, San Diego; Department of Neuroscience, University of Rochester School of Medicine and Dentistry; Center for Visual Science, University of Rochester School of Medicine and Dentistry; Intellectual Development and Disability Research Center, University of Rochester School of Medicine and Dentistry

## Abstract

Multi-electrode recording of neural activity in cultures offer opportunities for understanding how the structure of a network gives rise to function. Although it is hypothesized that network size is critical for determining the dynamics of activity, this relationship in human neural cultures remains largely unexplored. By applying new methods for analyzing neural activity to human iPSC derived cultures at either low-densities or high-densities, we uncovered the significant impacts that neuron number has on the individual neurophysiological properties of cells (such as firing rates), the collective behavior of the networks these cultures formed (as measured by entropy), and the relationship between the two. As a result, simply changing the densities of neurons generated dynamics and network behavior that differed not just in degree, but in kind. Beyond revealing the relationship between network structure and function, our findings provide a novel analytical framework to study diseases where network level activity is affected.

## INTRODUCTION

Neural networks differentiated from human induced pluripotent stem cells (iPSCs) (Aoi et al. 2008; Takahashi et al. 2007) have emerged as one of the most promising systems for modeling a vast array of human diseases ranging from neurodevelopmental disorders to neurodegeneration (Whiteley et al. 2022, Ardhanareeswaran et al. 2017, Vadodaria et al. 2020, Rivetti di Val Cervo et al. 2021, Wang et al. 2023). Neuronal cultures are especially compelling because they allow for study of disease pathology across multiple scales of complexity, such as gene expression, cellular physiology and synaptic transmission. Each level of analysis provides a different lens within which to view disease processes, and therefore identify mechanisms and potential areas of intervention. Interestingly, while the studies using human iPSC-derived neuronal cultures unravel the complexity and dysfunction of neural circuits at the molecular and cellular level are extensive (Chiaradia and Lancaster 2020, Kim et al. 2020 Eichmüller and Knoblich 2022, Andrews and Kriegstein 2022), approaches to study the emergent patterns of network activity are comparatively limited. Beyond the importance of studying neuronal activity to understand neural circuits as a self-contained goal, studying population activity serves as a critical bridge between the microscopic features of a circuit (the cells and the synapses) and the macroscopic behavior of the system as a whole. More broadly, population activity can provide critical links between the molecular underpinning of neurobiological diseases and change in human brain activity that can be measured by non-invasive methods, such as functional magnetic resonance imaging (fMRI) and electroencephalography (EEG). However, these methods lack the resolution of cell-to-cell interaction.

Functional assay of activity of neural networks in culture plates can be achieved using multi-electrode array (MEA), which is technology that allows recording of the extracellular voltage changes from neurons over time. Compared to traditional patch clamp, the MEA technology has the advantage of being high throughput; it has the capacity of recording electrophysiological activity of hundreds of neurons simultaneously over long periods of time. Remarkably the MEA data recordings of neural networks can be used as comparative functional readouts for different human neuronal states including healthy or diseased conditions and as a platform for drug discovery and drug testing (Hu et al. 2022, McCready et al. 2022, Pré et al. 2022, Vadodaria et al. 2021, Tukker et al. 2018, Pelkonen et al. 2022).

The most prosaic measure of a neuronal circuit is the number of neurons in the network. In the simplest sense, if a neuron (N) is either active or inactive at a moment in time, the number of combinatorial patterns that can be generated by a population of neurons is 2^N. Thus, an assumption of computing has been that the more neurons there are, the more patterns can be generated (Shannon 1948). This model however fails as neurons are connected to one another, and their activity therefore influences each other, thus violating a simple assumption of network size and pattern complexity, the independence of neuronal firing. As a consequence, a critical open question in systems neuroscience is the relationship between the size of the network, or the number of neurons in the network, and the computational complexity of that network. Answering such a question is central to understanding a number of features of neocortical organization, from the relationship of brain size to behavior across different species, to the impact that neuronal loss and neurodegeneration has on circuit function in neurodegenerative diseases.

To address the question of the relationship between the size of the network and its complexity, we measured neuronal activity of human iPSC-derived neural cultures of different sizes over 2 months using MEA. As the neurons matured, simple properties of neuronal activity such as firing rate increased in both low and high-density cultures, albeit with different time courses. Interestingly, when we examined the activity at the population level, we found that low-density cultures exhibited more complex activity patterns as compared to high-density cultures. Furthermore, we found that a statistical model which aims to predict the activity of the population at the macroscopic scale from the behavior of the microscopic components, was better at doing so for high-density cultures as compared to low-density cultures. Taken together, these data show that low-density cultures present more complex connectivity as compared to high-density cultures, suggesting that properties of neural network organization can vary in fundamental different ways simply by changing the density of the neurons in the network. Our results can have a potential impact on stem cell-based neurological disease modeling and the new analysis methods presented here can be invaluable for researchers studying *in vitro* human neuronal activity patterns.

## RESULTS

### Generation of iPSC-derived neural networks of different size

To determine the relationship between neuronal density and network organization, we cultured human iPSC-derived neurons for 8 weeks and recorded spontaneous extracellular activity weekly using MEA for 250 hours (N = 792 recordings) beginning at 3 weeks *in vitro* (WIV) and continuing until 8 weeks WIV. Cultures were plated at 5 different densities (2.5 K cells/cm^2^, 5 K cells/cm^2^, 12.5 K cells/cm^2^, 25 K cells/cm^2^, and 50 K cells/cm^2^, Figure. 1A). Cells were counted on the last day of recording and showed a strong correlation between the plating density and the final cell counts (Figure. 1B, Figure S1). Although, no significant differences were detected between the number of neurons (MAP2 positive cells, Figure 1C) or the number of glia (GFAP positive cells, Figure 1D) as assessed by immunohistochemistry across the different culture densities, we did find that cultures with densities higher than 10 K cells/cm^2^ had on average fewer glia than those with densities lower than 10 K. Interestingly, a number of recent studies suggest that molecular and cellular features of networks may be affected by cell density. For instance, cultures with 5K cells/mm^2^ have approximately 400 synapses/neurons while higher-density cultures, such as ones with 4500 cells/mm^2^ have far fewer synapses/neuron (150 per cell) (Cullen et al. 2010). There are no standards for determining culture density, and different labs use an array of different densities, suggesting that experimental variability in the initial conditions may impact the findings, and as a consequence the interpretation of network activity (Pré et al. 2022, Wen et al. 2022, Marchetto et al. 2019, Mok et al. 2022). Interestingly, many manufacturers of tissue culture reagents, suggest plating densities of 10K cells/cm^2,^; a density which provides a convenient delineation between cultures of low density and high-density cultures. In this framework, our experimental design allowed us to explore how the density of neuronal cultures impacted the activity patterns they could generate. If for instance, low-density (networks represented by the color green) and high-density networks (represented by the color purple) self-organized in similar ways, then we would expect no differences in either the activity of individual neurons or the joint activity of the population of neurons. If however, the density influenced the ways in which neurons connected to one-another, and thus influenced the emergent activity patterns, then a signature of these differences in network structure would exist in the functional measures of activity. To test these hypotheses, we used low-impedance 64 (8 × 8 grid) channel MEAs to study the spiking activity of the individual neurons in the neural population. Figure 2A shows an example of a recording of 8 out of 64 total channels that revealed the diversity of neuronal activity, including multiple single neurons (or units) in a given channel, and the electrical signature of a given unit across multiple channels. To isolate single unit activity, we developed an analysis pipeline (Figure S2 and Experimental Procedures) that allowed us to both identify the activity from multiple cells recorded on a single channel and the activity of a single neuron that was present across multiple channels. In this example, we identified 10 units across the channels such that some channels contained activity from only one unit (channel 4, 10, 26, 31, 48, and 62), while others (channel 57 and 64) contained spiking activity from multiple well isolated units (Figure 2A). We could detect well-identified units by plotting a low dimensional representation of the spike waveforms (Figure 2B). In each column (Figures 2B-D), the spikes from the units (Figure 2B, colored points) were plotted, along with random periods corresponding to noise (Figure 2B, black dots). The isolated neurons had waveforms (Figure 2C, colored traces), which were distinct from the noise (Figure 2C, gray traces). We further ensured that the data was from single units by examining the inter-spike interval (ISI) distribution. Units with ISI lower than 1.6 ms corresponding to the spiking refractory period were also eliminated, thereby further ensuring the quality of the recordings. Subsequently single-unit activity was plotted across time (Figure 2E) and as a raster plot, where each tick represents an action potential from the unit and space (Figure 2F) and where each colored circle represents the putative location of the unit in the culture. The location of the units was found by comparing the level of activity in the surrounding channels and triangulating the location of the neuron from these traces (see Experimental Procedures). Using this method of temporal-spatial fingerprinting (where the neuronal identity was determined by using both the temporal structure of the spiking waveform and the spatial map of electrical activity from each unit across the channels) we found that neuronal firing was highly varied both in the rates of activity, but also in the temporal structure of activity across units, including periods of quiescence and spiking (Figure 2E). Such an isolation of individual cell activity allows for the study of the microscopic electrical properties of individual neurons and the macroscopic properties of the collective dynamics of networks of different cell densities.

**Figure 1.**
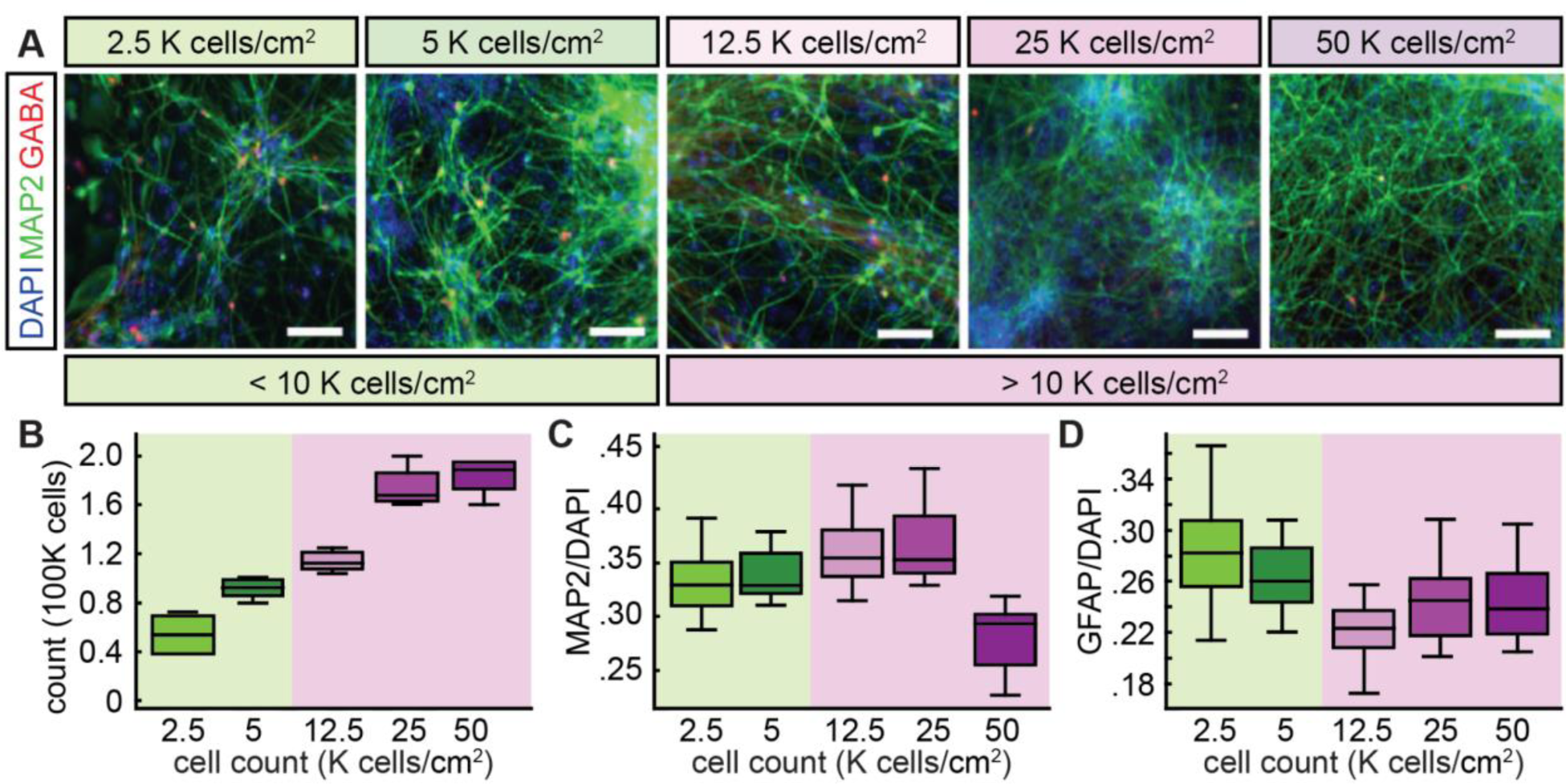
Generation of iPSC-derived neural networks of different sizes. (A) Representative images of immunostained cultures for the neuronal markers MAP2 (green) and GABA (red) at 8 weeks. Nuclei of all cells were counterstained with DAPI (blue). Scale bar, 100 μm. (B) Final number of cells at 8 weeks as a function of the initial number of plated cells (2.5, 5, 12.5, 25 and 50 K cells /cm^2^). Boxplot shows data from 4 wells. (C-D) Quantifications at 8 weeks of cells expressing the neuronal marker MAP2 (C) and the astrocytic markers GFAP (D) as a function of initial cell density. Boxplots show data from 7<n<12 randomized images.

**Figure 2.**
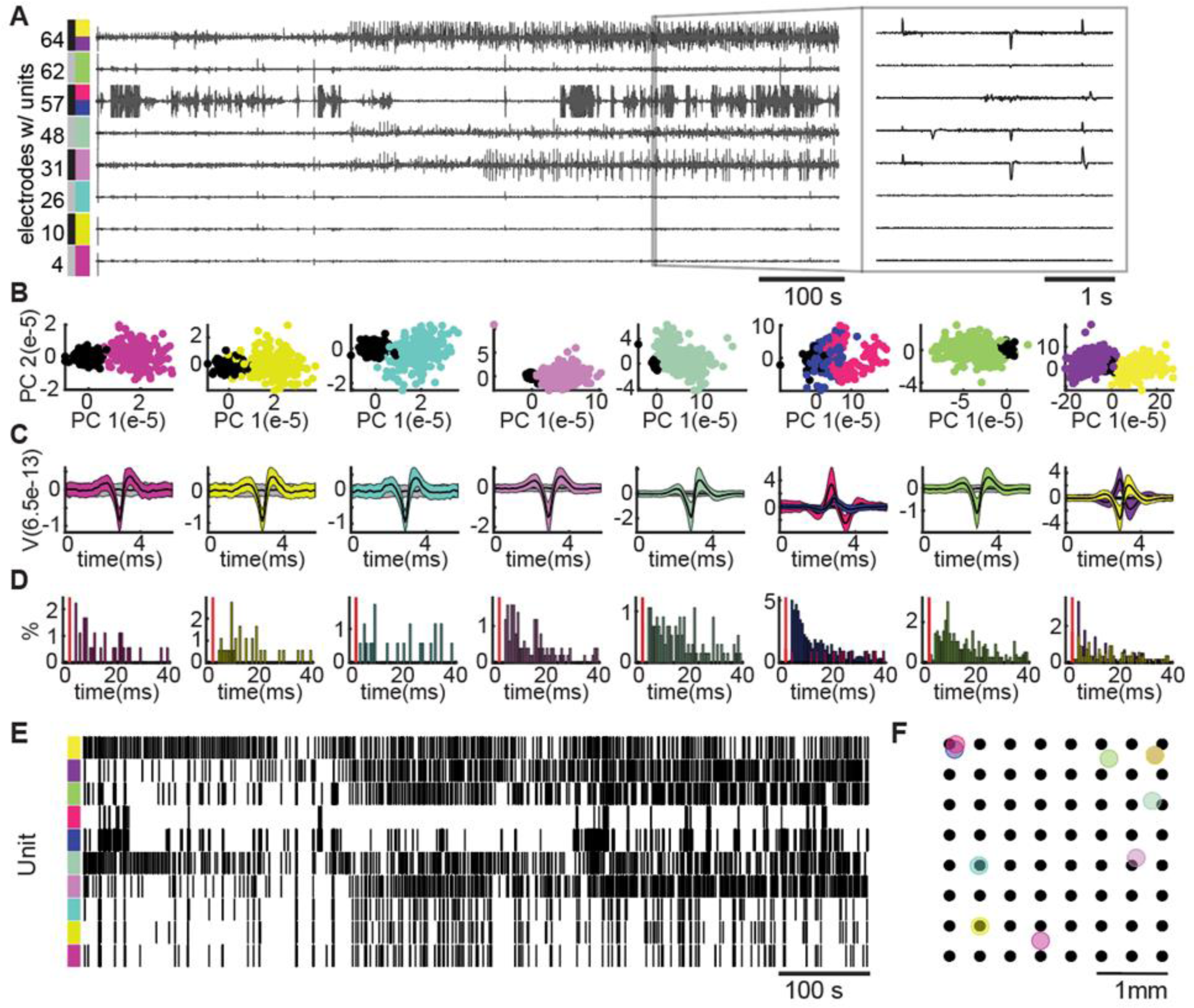
Spike sorting across multiple channels to isolate activity from single units. (A) Example of raw recording of 8 channels from the total 64 channels recorded in a single low-density culture showing clear spiking activity across the channels. Colors correspond to units identified following spike sorting in each channel. Insert shows the individual waveforms of the different units across time and across multiple channels. (B) Low dimensional projection of action potential (spike) waveform shape where each point corresponds to a single spike from a given unit. Colors correspond to the units and black dots represent random epochs from the recording corresponding to electrical noise. Waveforms from individual units cluster together in this space and are distinct from the noise. (C) Waveforms of the units detected (mean ± std) in B with colors corresponding to the isolated units and gray lines representing the noise from B. (D) Inter-spike interval (ISI) distributions for each unit with the red line corresponding to 5 ms. In addition to a clear non-symmetric action potential waveform, units were only included if there were at least 100 spikes that had an inter-spike interval of 5 ms or more. (E) Raster plot of the activity of the detected units from A following spike sorting. This allowed segregation of the activity from multiple units recorded on a single channel and to isolate the activity of a single unit that was recorded across multiple channels. (F) The putative spatial location of the unit was mapped by using a voltage telemetry approach to identify the location of the neuron within the 8 × 8 MEA.

### Microscopic features of network activity change with circuit maturation *in vitro*

To evaluate the statistical structure of the activity, we started by using the simplest feature of the detected units, the firing rate (Figure 3). We first observed that the neurons in low-density cultures had significantly higher firing rates as compared to the neurons in high-density cultures (Figure 3A). Although the high-density cultures had more neurons, the lower firing rates suggested that the structure of the network activity depended on more than just the neuronal number. We also observed that the average firing rates for the neurons in the populations increased from 3 to 8 WIV for both the low- and high-density cultures; firing rates varied over 1-2 log orders as neurons mature from 3 to 8 WIV (Figure 3B), indicating that neuronal maturation is taking place at both low- and high-density cultures, but at different rates.

**Figure 3.**
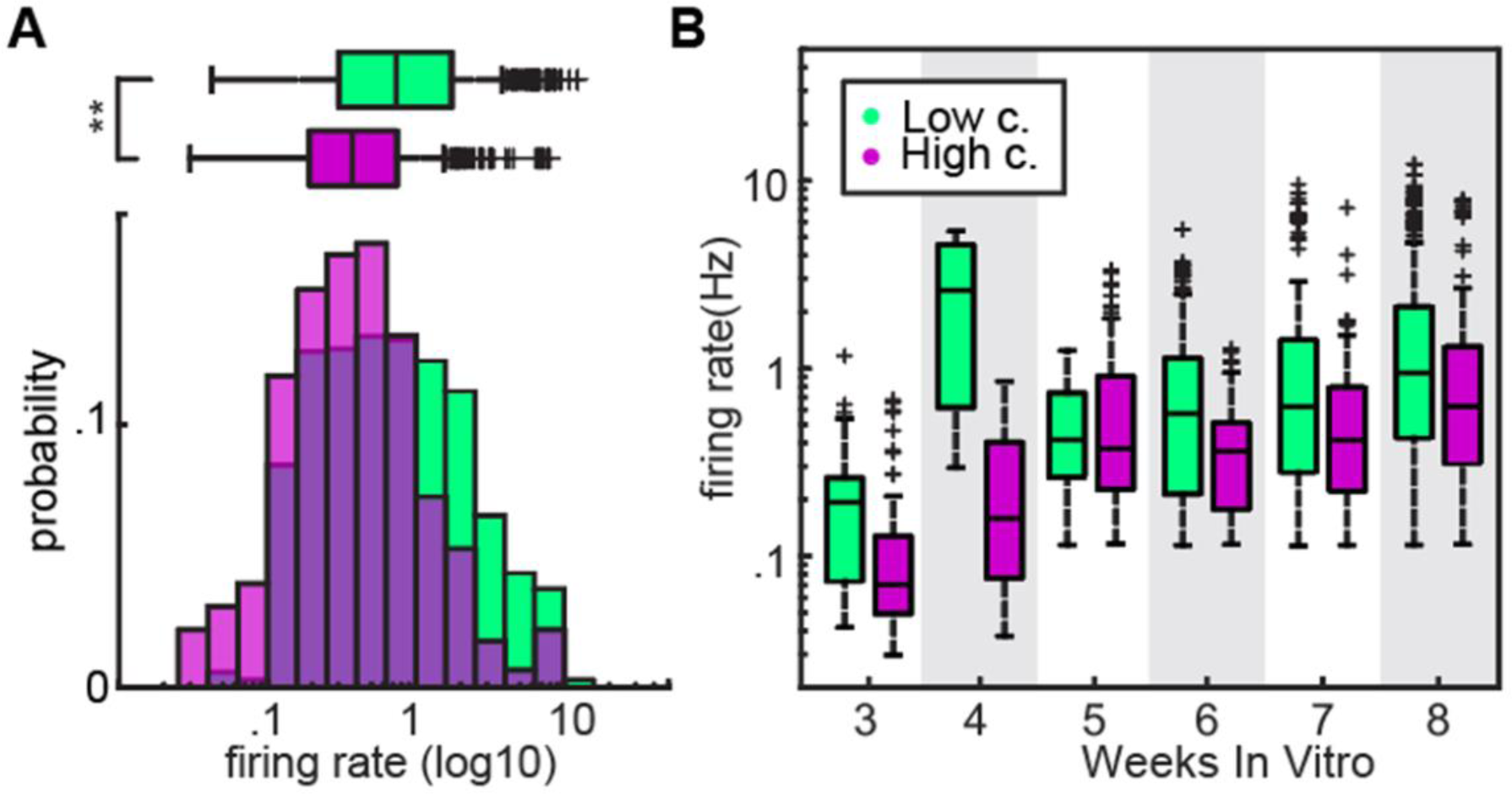
Firing rates of low-density networks are higher than in high-density networks. (A) The distribution of firing rates of low-(green) and high-density (purple) cultures is significantly different (low-density=958 units, high-density=434 units over all weeks *in vitro*, p< 0.0005 (P=4.4e-21). Statistics: two-sided Wilcoxon rank sum test. (B) Firing rates of low-density cultures and high-density cultures over the course of 8 weeks *in vitro* shows that low-density cultures had a significantly higher firing rate across time.

Other features of network activity can be quantified beyond the firing rates of individual neurons that capture aspects of the structure of neuronal circuits (Markram et al. 2004, Tremblay et al. 2016). For instance, all the cells in a culture may have similar firing rates, but the degree to which that firing is coordinated may vary substantially. Such differences in neuronal correlations can arise from differences in network connectivity, such as the synaptic organization of neurons in the population (Litwin-Kumar and Doiron 2012, Litwin-Kumar and Doiron 2014, Cohen and Kohn 2011). One way to evaluate this is to calculate the pairwise correlations of the individual units coming from the same recording. In Figures 4A-C, we show an example of a raster plot and correlation structure for both low-(Figures 4A1-C1) and high-density (Figures 4A2-C2) cultures. In this example raster plot of the low-density cultures (Figure 4A1) the activity patterns across time in different individual neurons are diverse. While a large group of cells fire synchronously across the population, there are numerous examples of cells that have temporal patterns of activity that are distinct from the overall population (Figure 4A). The pairwise correlations between neurons (Figure 4B1) reflect this difference and the distribution of these correlations across the population (Figure 4C1) quantifies this diversity in firing patterns. In the example of the low-density population, most of the correlation values are near zero, suggesting that the activity of many of the units is independent of one another (Figure 4C1). Importantly, we also see examples of correlation values ranging from near 0 to 1; a distribution that reflects different degrees of independence and synchrony across groups of neurons within the overall population. By contrast, for the example raster plot of the high-density cultures (Figure 4A2), activity patterns were not as diverse. To quantify this diversity in firing, we examined the distribution of pairwise correlations across low-density and high-density cultures. For high-density cultures, the mean correlation was close to 1, reflecting the hyper-synchronous activity we observed. By contrast, the correlation values in low-density cultures were much more varied, indicating that neuronal activity was much more complex (Figure 4D). Interestingly, the differences in correlation emerged in the earliest stages of culture maturation as measured by WIV. Low-density cultures started to show a higher correlation between pairs as early as week 4, whereas high-density cultures started to show this increase beginning at 6 WIV (Figure 4E). These data demonstrate not only that firing rates differ between low- and high-density networks, but that the interactions between neurons, as calculated by the pairwise correlations in the fluctuations of activity, also vary across cultures of low- and high density. This result implies the underlying organization (functional connectivity) between networks undergoes a change with an increase in network size.

**Figure 4.**
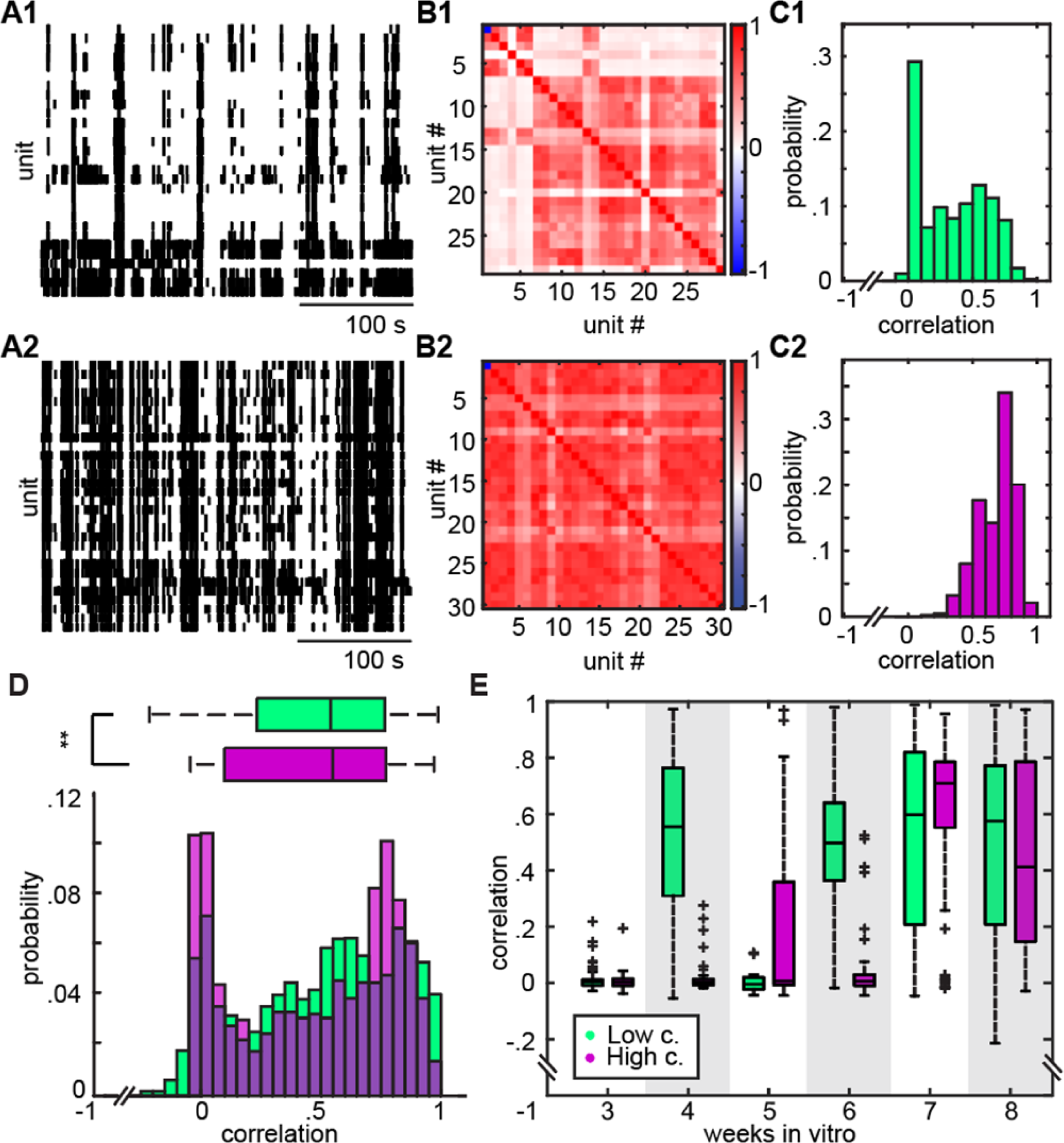
Differences in pairwise correlations reveal heterogeneities in functional connectivity across low- and high-density cultures. (A) Example of a raster plot for low-density (A1) and high-density (A2) cultures. (B) Pairwise correlations of the activities of the low-density culture in A1 (B1) and of the high-density culture in A2 (B2). (C) Histogram of the pairwise correlation values of the low-density culture from B1(C1) and of the high-density culture from B2 (C2). (D) A significant difference in the pairwise correlations from all recordings for low-density (green) versus high-density (purple) cultures reflects not only differences in the mean correlation but also in the distribution of correlations corresponding to large number of highly synchronous populations in high-density cultures (low-density = 6946 pairs, high-density = 1270 pairs, p< 0.05). Statistics: two-sided Wilcoxon rank sum test. (E) Pairwise correlations for low-density samples increased and diversified earlier, starting as early as week 4, and this diversity remained broader for the rest of the development compared to pairwise correlations of high-density samples.

### Macroscopic organization of population activity differs with network density

Measures of population activity such as firing rate or the pairwise correlations provide what one might think of as microscopic details about the iPSC-derived neuronal network, the way that one element or pairs interact. Next, we wanted to examine the macroscopic or global behavior of the populations of neurons in the cultures. A number of methods have been developed in other systems, including *in vivo* recordings in awake behaving animals and humans to quantify the pattern of activity across populations of neurons (de Ruyter van Steveninck et al. 1997, Dayan and Abbott 2005, Averbeck et al. 2006, Williamson et al. 2019, Vyas et al. 2020). The most straightforward of these methods is entropy, which describes the distribution of activity patterns across the populations over all time. Consider, for example, a network with highly synchronous neurons that all fire together. At any given time, the network will be in one of two states, either all the cells will be quiescent, or all the cells will spike. This would be an example of a low entropy network, with only two states. By contrast, where every combination of activity patterns across the population is equally likely, the entropy would be high. In practice, networks are rarely in these two extreme states, but they illustrate one approach by which the overall diversity of activity patterns can be statistically described and shed light on how this activity may be related to network structure.

To quantify the entropy of low- and high-density cultures, time series were binarized considering 11 ms bins of time, such that if a spike occurred within this window, it was registered a value of 1 and if no spike occurred, it was registered as a 0 (Figure 5A). This bin size ensured that over 95% of the 11 ms windows where action potentials were detected had only one event in the bin (Figure S3). In this framework, two features of the network activity can be quantified, the count entropy (how many neurons fire at the same time) and the word entropy (the pattern of neurons that fire within a given time window). Count entropy is most similar to the burst measures of neuronal activity in human cultures that have been used previously and serve as a proxy for population synchronization (Maeda et al. 1995). By contrast, the word entropy, which measures not only how many cells are active within a burst, but also which cells are active, is more sensitive to the identity and nuances of patterns of firing, and thus provides a richer portrait of the activity in cultures (Osborne et al. 2008). In this example, the probability distribution of counts (Figure 5B) and of words (Figure 5C) show that quantifying networks in terms of word patterns is necessarily richer in terms of providing information about the network. Indeed, when we calculated the count entropy versus the word entropy for this example (Figure 5D) and across all experiments (Figure 5E), we found that word entropy was always significantly higher than count entropy in bits and this held across networks with different numbers of neurons (sampling size) (Figures 5D-E). Given this richness, we quantified the word entropy (which we shall call simply entropy) for low- and high-density cultures and found rather surprisingly that the lower density networks had a significantly higher entropy as compared to the higher density networks (Figure 5F). This result held across networks with different neuronal numbers (Figure 5G), further suggesting that the overall architecture of low- and high-density networks was different. Interestingly, when we examined how the entropy changed as a function of time, we found that within 4 weeks, low-density network entropy was significantly higher than that observed in high-density networks (Figure 5H). Although the entropy for the higher density networks increased with maturation time, the diversity of patterns generated by lower density networks remained significantly higher across all times. Taken together, these data suggest that both the complexity of patterns generated, and the way in which that complexity emerges as networks self-organize *in vitro* differ as a function of network density. Rather counterintuitively, we found that lower density networks, which one would predict could carry less information, actually produced more complex activity patterns than higher density networks.

**Figure 5.**
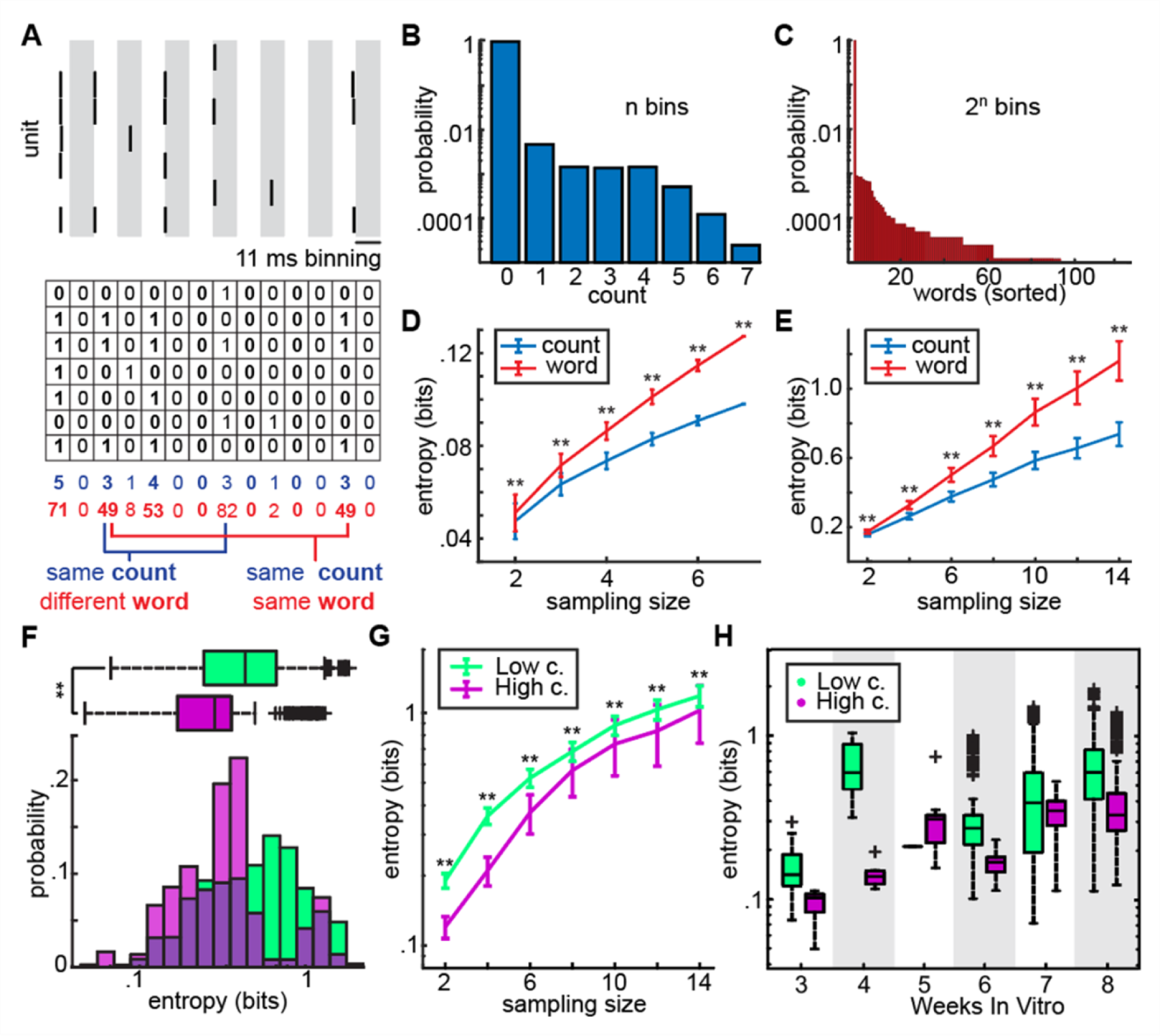
Characterization of population activity using entropy reveals differences between low- and high-density cultures. (A) On top is an example of spiking across a population of 7 neurons. The activity is binned into 11 ms intervals (interleaved with gray shading to show bins). On the bottom, the spikes were binarized into patterns of 1 s (1 or more action potentials within a bin) and 0 s (no action potentials in a given bin). For each 11 ms window was calculated the count (blue) corresponding to the number of units active and the word (red) corresponding to the pattern of neuronal activation. For 7 units, there can be 8 different patterns of activity when measuring counts and 128 different patterns of activity when measuring words. (B) Distribution of counts for the population in A across all time represents the probability of seeing any given pattern of activity. The distribution of counts corresponds most similarly to measures of synchrony and bursting. (C) Distribution of words for the population in A across all recording time shows the likelihood of seeing any combination/pattern of neuronal activity. (D) Word entropy (red) is higher than count entropy count entropy (blue) across all population size (different numbers of neurons) showing that word entropy captures more of the richness of population activity than count entropy (100 subsamples of this neuronal population (N=2-7 neurons) where a total of 7 neurons were recorded. (p < 0.005 for all samples with multiple pairwise comparison correction; p = 2.6e-4, 8.3e-22, 3.4e-33, 2.4e-34, 1.9 e-34, 9.6e-41 for k = 2,3,4,5,6,7 respectively). (E) Word entropy is higher than count entropy for all recordings across different numbers of neuronal populations. For each sampling size (count and word resamples for different numbers of neuronal populations defined by K: K(2)= 3172, K(3) = 3056, 2404, 2001, 1569, 1415, 1071, p<0.005 with correction for multiple pairwise comparisons). (F) Low- and high-density cultures have significantly different word entropy distributions (low-density=1962 populations, high-density=362 populations, p<0.0005). (G) Low-density cultures have significantly more entropy than high-density cultures across populations with different numbers of neurons/population (low-density populations = 2411, 2382, 2042, 1753, 1368, 1215, 920, high-density populations = 761, 674, 362, 248, 201, 200, 151, p<0.005 across all population with correction, p= 4.4e-44, 1.7e-65, 1.5e-22, 2.4e-5, 3.2e-5, 4.9e-8, 9.5e-4 for neuronal populations = 2, 4, 6, 8, 10, 12, 14 respectively). (H) Comparison of word entropy distribution across weeks *in vitro* for low-(green) and high-density (purple) cultures.

Our analyses thus far on the firing rates, the pairwise correlations, and the overall network activity patterns reflect two different scales of analyses. At the microscopic scale, we see a difference in the behavior of individual neurons and differences in their pairwise functional coupling. At the macroscopic scale, we see differences in the overall organization of activity across ensembles of neurons in the network. A critical question we can ask is how the microscopic features of neuronal networks determine the macroscopic behavior of the culture as a system. One approach to do this is to use an Ising or maximum entropy model from statistical physics (Jaynes 1957, Schneidman et al. 2006). These predictive models have been used to understand how complex behavior in a system arises from the behavior of the elements of the system, such as for neurons in the retina (Shlens et al. 2006), olfactory bulb (Chockanathan et al. 2021), or hippocampus (Meshulam et al. 2017, Chockanathan et al. 2020, Chockanathan and Padmanabhan 2021). In a pairwise model, the goal is to predict the empirical data (the distribution of activity patterns that arise at the level of the population) from the behavior of the elements (the firing rate of the neurons and the pairwise correlations of their activity) with as few assumptions as to what that structure is (Jaynes 1957a) (Figure 6A). To do this, we first subsampled different units in the detected population, for instance 6 random units from a population of 10 neurons (Figure 6A, left). The activity of this population (the spiking patterns) is divided into two parts, one that is used to generate distribution from a predictive model, and the other that will be used to assess the goodness of these predictions. The predictive model learns the features of spiking and correlation and optimizes the parameters h_i_ (neural excitability parameter) and j_ij_ (neural interaction parameter), respectively, to make predictions about what the overall activity patterns will look like (Figure 6A, middle). A comparison of low- and high-density culture samples is given in Figure 6B for the pairwise model. The close the points are to the unity line, the better the prediction of the model. One can observe that the model has a relatively poorer result for the low-density sample on the top, as the distribution is more spread around the unity line. We then used a measure called the Kullback-Leibler Divergence (KL-D) which quantifies how much the predicted probabilities of different patterns deviate from the observed patterns in the data (Figure 6A, right). The higher the KL-D, the poorer the model is at predicting the data. First, across all experiments, we found that the second-order model was significantly poorer at predicting the structure of overall activity in low-density cultures as compared to high-density cultures (Figure 6C). Second, we observed that the second-order model was poorer at predicting the network structure of the low-density cultures early in development and remained so up until 7 WIV (Figure 6D). These data suggest two important aspects of network self-organization. First, the patterns in low-density populations exhibited a structure that was sufficiently more complex than that of high-density populations. Second, this complexity appeared to emerge early in the self-organization of the networks (Figure 6D). To ensure that these models accurately recapitulated the elements of the microscopic behavior, we compared the h and J terms from the model across the two culture densities. As with firing rate, we found that the h term which corresponds to excitability was significantly lower in high-density cultures as compared to low-density cultures (Figure 6E, top). Furthermore, we found that the interaction term, J from the model, was also significantly different between low- and high-density cultures, recapitulating the differences we saw in the pairwise correlations we calculated (Figure 6E, bottom).

**Figure 6.**
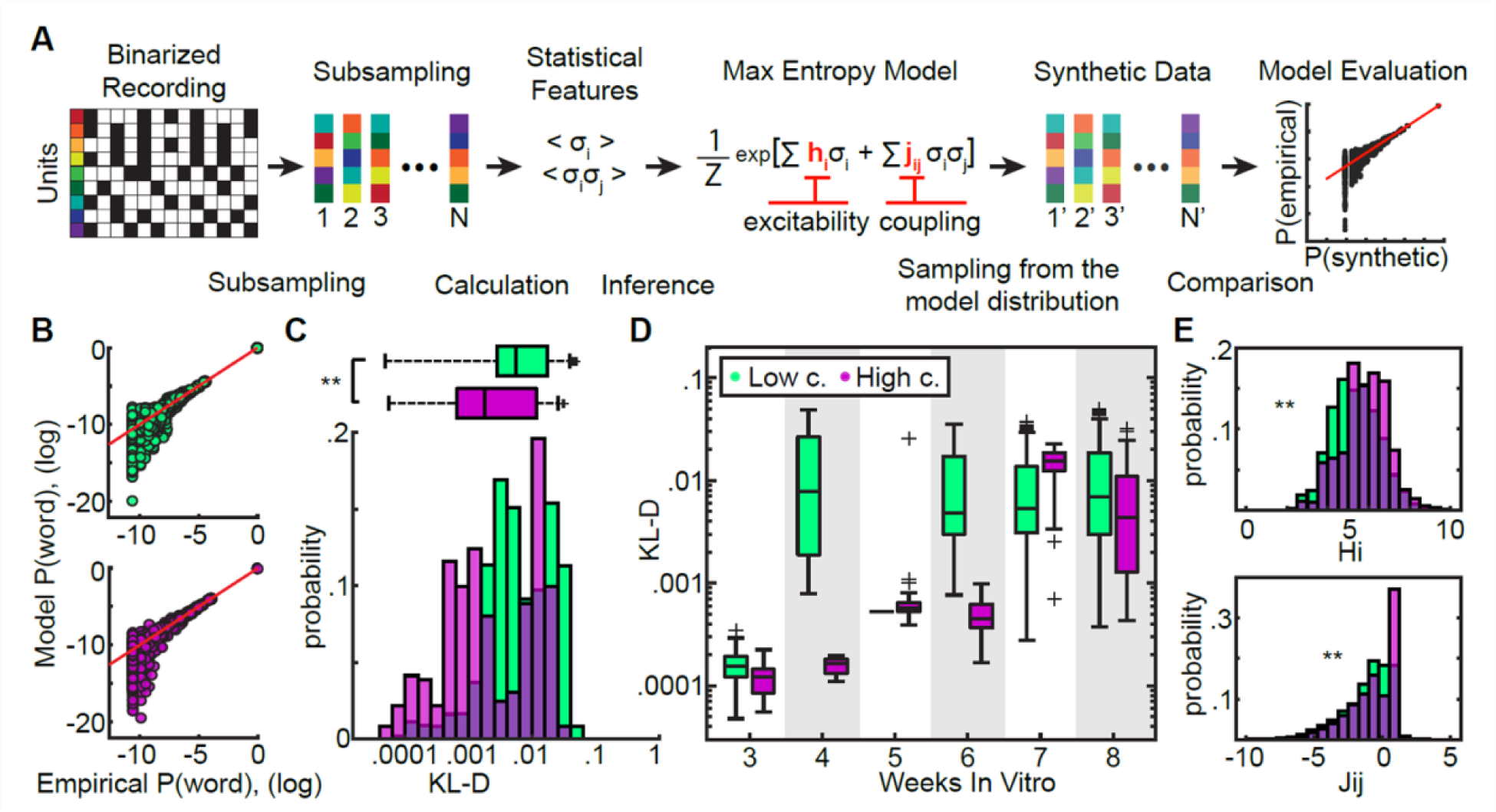
Maximum-entropy models link the firing rates and pairwise interactions between the individual units to the global activity of the population. (A) Schematic representation of the maximum entropy model analysis from recorded data to model generation and fitting. First, the neural activity is binarized. Following this, we randomly selected subgroups of neurons from the total population of neurons that were recorded. The statistical features of the population activity (firing rate) and the pairwise interactions were extracted for each random subgroup and a maximum entropy model was used to create synthetic data about the distribution of global patterns of activity across the population. This synthetic data was then compared to the empirical data to determine the goodness of fit of the model. (B) Example of a pairwise maximum entropy fit for a low-density (green) and high-density (purple) culture. Each point corresponds to a single pattern of activity in the population and the red line is the unit line. The pairwise model is less successful at predicting the different patterns of activity observed in the low-density sample as compared to the high-density culture. (C) Low- and high-density cultures have significantly different distributions for Kullback-Leibler Divergence (KL-Divergence) (n_low = 1962, n_high = 362 KL-D values, p = 1.0e-25). (D) KL-Divergence across weeks *in vitro* show that second order models of neuronal activity are worse at predicting the structure of the populations in low-density cultures as early as 4 weeks *in vitro*. These differences show that low-density cultures exhibit more complex patterns of activity earlier in vitro as compared to high-density cultures. (E) Low- and high-density cultures have significantly different distributions for model parameters of firing rate (Top) (low = 11772, high = 2172 H_i parameter, p = 2.6e-59) and pairwise interactions (Bottom) (low = 29430, high = 5430 J_ij parameter, p = 2.7e-66).

Taken together these data reveal that simply changing the density of the neurons in the neural network has a significant impact on the behaviors of the individual elements (the neurons), the collective behavior of the system (the entropy) and the relationship between the parts and the whole (the pairwise maximum entropy/ Ising model).

### Low-density networks show organization across different spatial scales

We finally wanted to assess what the correlations, the entropy, and the maximum entropy fit for network activity might reveal about the functional coupling and the organization of the cultures *in vitro*. To do this, we returned to the spatial maps of neurons in the 8 × 8 recording array. As the biophysical fingerprinting (Figures 2 and S2) provided a location in space of the recorded units, we wished to connect the physical distribution of the neurons to their physiological measures (activity and pairwise connectivity) (Figures 7A1-2). In this representation, each neuron is a single color with its center corresponding to the physical location of the cell in the culture and the gray line connecting any two neurons reflects the correlation in the time series of activity. Here, the radius of the circle represents the firing rate of the neuron. Additionally, the thickness of the gray line corresponds to the correlation value. As any two time series will produce some correlation, we only included connections that were above a threshold of 0.2 consistent with previous studies estimating functional connectivity (Chockanathan and Padmanabhan 2021). When we examined connections after a threshold was applied, we first noticed that high-density cultures appeared to have connections across multiple cells distributed throughout the culture. By contrast, connections of the low-density cultures were more spatially close together. To represent this, we examined the distance d across all pairs and plotted the distribution of this distance (Figure 7B), and its relationship with the distribution of pairwise correlations for low- and high-density samples (Figures 7C1,C2 respectively). First, we noticed that units from high-density cultures were not connected to every other cell in the population. Connections were fewer, but the activity was highly synchronous across the populations. In low-density populations by contrast, the network appeared to be connected more densely, but with connections weaker to the overall population. Second, we observed a diverse physical arrangement of functionally connected neurons in low-density cultures (Figures 7A1,C1). For example, for a given correlation, we found examples of pairs that were both spatially close and spatially far and this difference was observed over an order of magnitude (from 1mm to 100 um). By contrast, in high-density cultures, connections across all spatial scales were high. (confined to the upper band (Figures 7A2,C2), suggesting the functional connectivity was homogeneous. Taken together, our results suggest that the smaller coding capacity and dynamics of high-density networks arises from the spatial homogeneity of their network organization.

**Figure 7.**
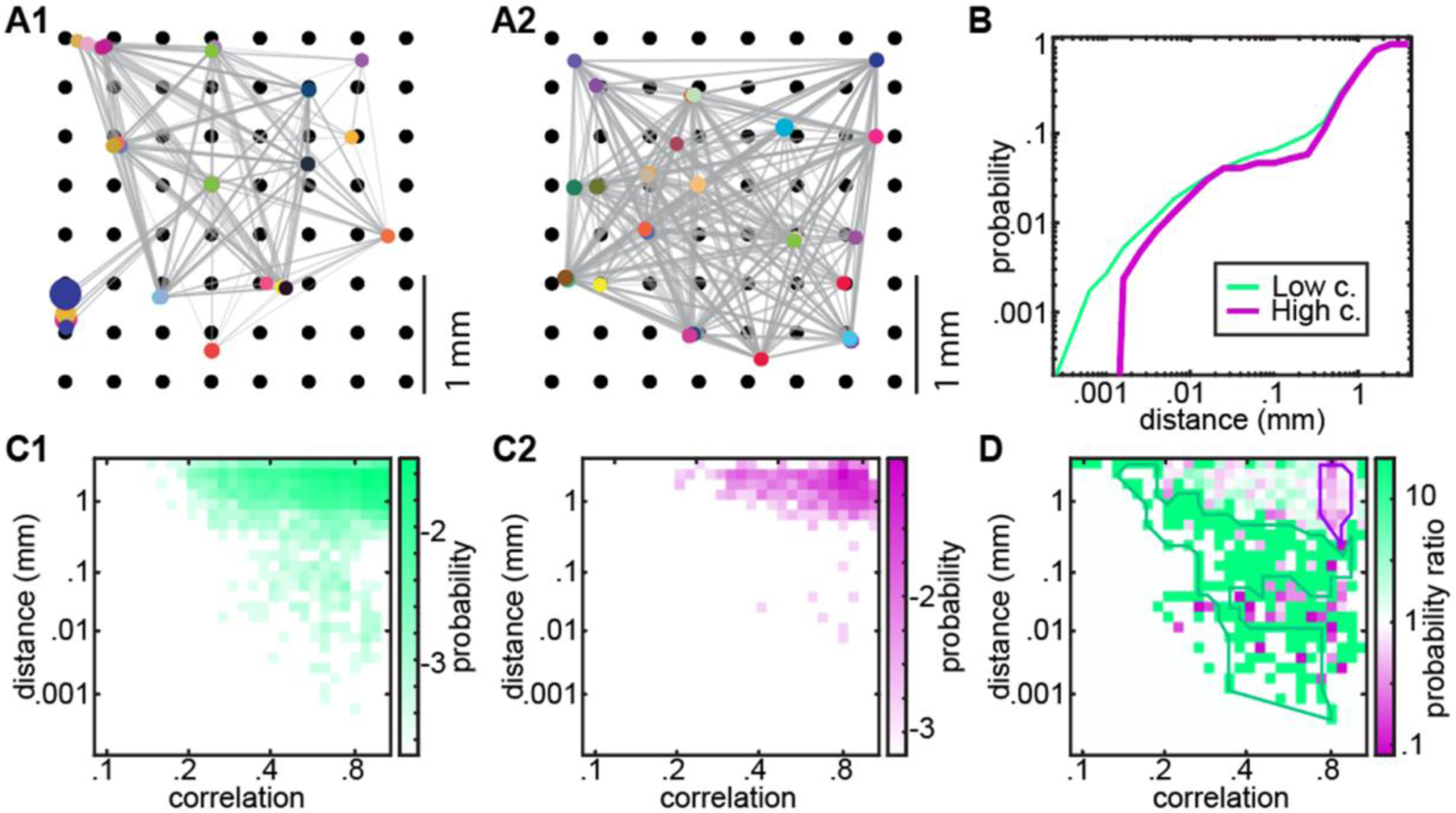
Low-density cultures show more complex patterns of functional connectivity across different spatial scales. (A) Example of functional connectivity detected by 8×8 Multi-electrode array (MEA) for a low-density culture (A1) and a high-density culture (A2). Black circles correspond to the physical location of channels). Each floored circle represents a unit and its size is proportional to the firing rate. Each gray line reflects a pair-wise correlation above a significant threshold between the two connected units. The gray line thickness is proportional to the correlation. (B) The physical distance between functionally connected units for low-density (green) cultures is more diverse than the ones for high-density (purple) cultures (low = 5210, high = 860 distance values, p = 0.04). (C) Probability distribution in correlation-distance space for low-density cultures (C1) and high-density cultures (C2). (D) Probability ratios of low- and high-density cultures emphasize the difference between the distributions. Low-density cultures occupy a wider range in both correlation (how variable are functional connections) and distance (how variable is the spacing of neurons from one-another). The higher probability ratio across multiple spatial and functional connectivity scales in low-density cultures reveals that networks of low-density are organized in more complex ways as compared to high-density cultures.

## DISCUSSION

Human monolayer cultures are a powerful tool for modeling both normal human neuronal networks as well as those disrupted in various disease processes. By varying one of the parameters of the culture, the density of neurons, we found a number of aspects of self-organization and functional coupling were revealed from the circuit’s activity.

Our observation that mean firing rates were higher in lower density cultures as compared to high-density cultures suggests that *in vitro* human neuronal cultures could be scaling mean firing rate based on principles of homeostasis. Cannon was perhaps the one of the earliest to appreciate the tension between the unstable and changing structures that compose biology and the stable function that emerges from that biology (Cannon 1932). More recent experimental and theoretical threads to unravel the neurobiological mechanisms of network stability and homeostasis (Marder and Prinz, Turrigiano and Nelson, and others), point to compensatory mechanisms such as synaptic scaling and firing rate normalization (Marder and Prinz 2002), often in the context of either *in vivo* development (Turrigiano and Nelson 2004, Garcia-Bereguiain et al. 2016), or *in vitro* perturbations to network activity (Turrigiano et al. 1998, Burrone et al. 2002). Our work here suggests that even something like neuronal density may exert strong effects on firing rate. Although we cannot measure synaptic strength using extracellular MEAs, our data suggest that the lower firing rates of high-density cultures may correspond to increased connectivity between neurons that are tightly packed together. Indeed, when we measure the pairwise correlations between neurons in higher-density cultures, they are greater than those observed in low-density cultures, suggesting that these high-density networks are more functionally connected. When we examine what kinds of patterns can be generated by these *in vitro* networks, we observe that high-density populations spontaneously generate fewer and more stereotypic patterns of global network activity as compared to networks of lower density. Although compelling, this link between cell density and entropy (the coding capacity of the network) does risk falling into a tautological trap. Higher correlations necessarily mean more similar patterns of activity, which by construction reduces the coding capacity. To unpack if our lower entropy result in high-density cultures was a trivial consequence of either the correlation or the firing rate, we turned to maximum entropy (Ising 1925; Jaynes 1957b) models that were originally developed in the field of statistical physics to determine how much of the complex global structure of a system could be described by knowing the microscopic interactions that were occurring across the parts of the system. In its simplest form, the pairwise maximum entropy model determines how well one can predict the distribution of the global states of the system (in our case, the patterns of activity of the whole network) knowing only the firing rates and the pairwise correlations. The KLD, a measure of the accuracy of the prediction, reveals how much extra information is stored in the activity of the system beyond the structure of the parts (Kullback and Leibler 1951). In this way, were the global differences in patterns of activity between low-density and high-density cultures only due to the correlations and firing rate differences, we would see no differences in the KLD values for low-versus high-density cultures. Surprisingly, we found that the KLD for low-density cultures was higher than that of high-density cultures revealing that the former self-organize in ways that are more complex, and therefore carry more computational capacity than cultures where the density of neurons is higher. An analysis of the spatio-temporal organization of the network appears to bear this out. Whereas the structure of high-density networks are all confined to narrow bands of space and correlation, the connectivity of low-density networks span many log-order scales of space and correlation.

A nt very low densities, neural cultures either do not survive (failing to form connections between neurons that ultimately leads to neuronal death), or are arrested as early differentiation states wherein a complex network structure is absent. We find that beyond this critical lower bound, the relationship between the density of neurons in the culture, and the organization of the emergent activity in that culture is not simply monotonic (more neurons = more complexity), but instead shows a complex relationship between network size, network connectivity, and computational capacity. The most parsimonious description of this would be that some Goldilocks size exists that maximizes coding capacity based on network density. Such relationships have been observed in other complex self-organizing systems including across networks that are varied as quantum cellular automata (Hillberry et al. 2021) and food webs (Petchey et al. 2008). This claim must be tempered by a number of specifics to neuronal circuits. First, our networks self-organize, and it is unclear if these different circuit principles would be preserved if the networks were assembled during development within the three dimensional structure of the nascent nervous system where signaling cues that guide the formation of that system are critical for synaptic connectivity and circuit development. Additionally, much of development is influenced and instructed by patterns of activity that arise from external inputs, most often via sensory experience. The extent to which sensory experience or developmental cues either make networks of different densities more similar or different cannot be answered by our study.

Furthermore, the coding capacity benefits of low-density networks must be considered within the context that no particular computation is being optimized or performed by these networks. One could easily imagine that the different network densities, and the resultant dynamics seen in our in vitro experiments, recapitulate the different computations that a complex nervous system, with a diverse array of circuits, performs. The hyper-synchronous activity seen in high-density connected networks may be more reflective of circuits where a single dynamic behavior, for example rhythmic firing, is required for a function. Examples of this highly stereotyped correlated activity exist throughout the nervous system and include constricting an abdominal ganglion (as in the lobster) or in mammal locomotion (Hudson et al. 2010, Hallinen et al. 2021). By contrast, a low-density network, with connectivity that spans multiple spatial scales, may more faithfully recapitulate the organization of a circuit in the associative region of the brain, where an increased coding capacity is essential for integrating diverse sensory and motor information to flexibly execute behaviors.

The considerations notwithstanding, our analyses reveal a framework for relating the elemental of a neuronal circuit, for instance the firing rate and the pairwise correlations between active neurons to the global structure and patterns of activity throughout the brain. Maximum entropy models, like the ones we have applied to *in vitro* human neuronal cultures have been used to study the emergence of structure throughout the nervous system as well as in modeling a number of diseases (Ashourvan et al. 2021, Nghiem et al. 2018, Chockanathan et al. 2020, Chockanathan and Padmanabhan 2021). In this regard, we present here not only a result about the relationship between the density of neurons in a network and the structure of activity that can be generated, but a way of investigating the global structure of activity more broadly in a number of applications related to human neural cultures ranging from development to the neurobiology of brain disorders.

## EXPERIMENTAL PROCEDURES

### Resource availability

#### Corresponding authors

Krishnan Padmanabhan and Maria C. Marchetto

#### Materials availability

This study did not generate new unique reagents.

#### Data and code availability

Raw data and code are available upon request to

### iPSC lines and neuronal differentiation

The two iPSC lines used in this study were derived from healthy controls, one female and one male, as previously described (Marchetto et al. 2010): WT-33 (RRID:CVCL_HA45) and WT-126 (RRID:CVCL_YL15). Samples were obtained with approval from the internal review board of the The Salk Institute for Biological Studies and informed consent of all subjects. Neurons were differentiated from iPSCs through intermediate generation of neural progenitor cells (NPCs) as described (Marchetto et al. 2019). Briefly, iPSC colonies were dissociated from Matrigel (Corning)-coated plates with 1 mg/ml collagenase IV (Thermo Fisher Scientific) and plated onto low-adherence plates in mTerR1 medium (STEMCELL) to form a suspension of embryoid bodies (EBs). The day after, the medium was changed to NPC medium: DMEM/F12 medium with Glutamax with 1 × N2 supplement and 1 × B27 supplement (all from Thermo Fisher Scientific) plus 500 ng/ml recombinant human noggin (Proteintech). After 10 days in suspension, the EBs were plated onto 10 μg/ml poly-ornithine (PLO, Sigma Aldrich) and 5 μg/ml laminin (Invitrogen) coated plates and maintained in NPC medium plus 500 ng/ml noggin and 1 μg/ml laminin. The neural rosettes were collected after 7 days, gently dissociated with accutase (STEMCELL) and plated in PLO-laminin-coated dishes. This NPC population was expanded using the NPC medium plus 10 ng/ml FGF2 (Joint Protein Central) and 1 μg/ml laminin. To differentiate the NPCs into neurons, the cells were cultured in DMEM/F12 Glutamax supplemented with 1 × N2 and 1 × B27, 1 μg/ml laminin, 20 ng/ml BDNF (Prepotech), 20 ng/ml GDNF (Prepotech), 0.2 μM ascorbic acid (STEMCELL), 1 μM retinoic acid (STEMCELL) and 500 mg/ml cyclic AMP (Tocris), with medium changes three times a week.

### Immunocytochemistry

Cells were fixed in 4% paraformaldehyde (Sigma Aldrich) for 10 min at room temperature. Antigen blocking and cell permeabilization were done using 5% horse serum and 0.3% Triton X-100 in PBS for 20 min at room temperature. Primary antibodies prepared in 5% horse serum were incubated overnight at 4°C, washed with PBS and incubated with secondary antibodies (1:500, Jackson Laboratories) for 1 h at room temperature. The primary antibodies used were the chicken anti-MAP2 polyclonal (1:1000, Abcam 5392), and rabbit anti-GFAP polyclonal (1:5000, Thermo Fisher Scientific 01670276). Cells were counterstained with DAPI for cell nuclei visualization.

### MEA recordings

12-well MEA plates from Axion Biosystems (San Francisco, CA, USA) were used to record electrical activity of neurons. The NPCs were plated in tetraplicate in previously PLO-laminin-coated plates at different densities and neuronal differentiation started the day after. Cells were fed three times a week and measurements were taken before feeding. Extracellular recordings were performed in a Maestro MEA system and AxIS software (Axion Biosystems) with voltages recorded at a frequency of 12.5 kHz. Following 5 min of plate rest time, recordings were performed for 15 min.

### Spike sorting using spatial information

#### Detection of spikes

To focus on the frequency region of spikes, the bandpass filter was applied to the raw signal recording between 500 Hz and 3500 Hz using the ‘butter’ function of MATLAB. Four-second padding was applied to the recording during the filtering process to prevent edge effects and paddings were removed afterward. After the bandpass process, data points that are at least 5 standard deviations away from the mean of the recording of each channel were detected as possible spiking points. From these points, only the ones that are maximum in their neighborhood (± 4ms) were accepted as spikes. This rule prevents double counting and neglects a spike occurring within the refractory period of another spike. The length of the waveforms was assigned as 8 ms for this step and for the pre-processing step. An upper threshold of 10 standard deviations was set to prevent the detection of noise and recording artifacts as spikes.

#### Pre-processing

To validate the spikes detected, they were tested against biological arguments. First, waveforms were trimmed from the ends and only the center 4 ms of the waveforms were taken for pre-processing. All spikes were scaled by dividing them by the absolute maximum value of their waveform. Afterward, the elimination process was applied using biological arguments, no symmetric waveform, and no close peak to trough. Since a spike waveform is a combination of two consecutive peaks with opposite signs, we expected to detect an asymmetric waveform. Accordingly, no symmetric waveform argument eliminates the spikes with similar first and second halves. For each spike, the dimension of the second half of the waveform was flipped, and the correlation and difference of both halves were calculated afterward. If the correlation was greater than 0.4 and the absolute value of the difference between the halves was smaller than 8, the spike was eliminated. Additionally, a sudden change in the raw recording due to a disturbance to the recording system might be detected as a spike after the bandpass process. No close peak-trough argument eliminates these noise spikes. Even if these noise spikes have an asymmetric shape, the distance between the peak and trough of the waveform of these spikes is quite narrow. Spike waveforms with peak-trough distances smaller than 0.32 ms were eliminated at this step. Parameter values were optimized with trial and error, according to their effect on the spike sorting procedure. Parameter assumed to be better if it results in better separated units and more well-defined waveform.

#### Sorting

The sorting algorithm we wrote facilitates the information coming from multiple channels at the same time. At every iteration, it focused on 4 channels of the 8 x 8 MEA. These 4 channels sit at the corner of a square e.g., the first 2 channels of rows 1 and 2. These 4 channels are the most informative channels for the region within the square they constitute. With this assumption, we iterated our sorting algorithm over the square regions over the MEA instead of iterating over the channels one by one i.e., we iterated the algorithm, not 8×8=64 times but 4×4=16 times. At each iteration, we collected spike times from all four channels and generated a new concatenated waveform for this group of channels (Gray et al. 1995). During the concatenation process, we only took the central 5.6 ms of the waveforms to concentrate on the part of the waveform that helps to differentiate the units better. The resulting waveforms were 4 x 5.6 ms long. Afterward, these new waveforms are sorted instead of the original waveforms. The advantage of using concatenated waveforms is that they can differentiate the physically separated units even if the units have a similar waveform. Also, a unit detected at multiple channels will be captured as a single unit within this square region.

Important features of the waveforms are extracted by projecting them over the first three eigenvectors of their covariance matrix (principle components). This lower-dimensional representation of the waveforms was clustered with the ‘evalclusters’ function of MATLAB. We chose the Gaussian mixture clustering algorithm as the clustering method and ‘CalinskiHarabasz’ as the evaluation criterion. We allowed the function to find up to 10 different units for these four channels. If the waveforms of different clusters had a correlation greater than 0.8, they were grouped together after clustering. Correlation values were calculated with the ‘corrcoef’ function of MATLAB. The resulting units were assigned to the channel for which the mean waveform has the highest amplitude. This procedure is the extension of the spike sorting procedure on single electrodes described by (Lewicki 1998).

Before moving to the next four channels, we inspected the units detected at the boundary between the current square and the next square. With the spike times of a given unit, the unit waveform was recreated using the recordings of the neighboring channels of the next square. Then, the amplitudes of the waveforms are compared. If the unit had the highest amplitude in the channel it is assigned to in the first place, nothing would be done. Otherwise, it is deleted and left to be sorted in the next square. This consecutive sorting and comparing process was repeated until we reached the opposite corner of the MEA.

#### Post-processing

During spike sorting, we combined the spike times coming from different channels. However, the same event might have been detected in different parts of the waveform at different channels. It is important since it can contaminate the inter-spike interval (ISI) distribution. For every unit detected, we removed the events which did not have their maximum at the center 2.4 ms region of their waveform. The waveforms of the remaining events are centered around their maximum value. Correlation coefficients between these centered waveforms were calculated with the ‘corrcoef’ function of MATLAB, and the duplicate waveforms with correlation value one were eliminated. In the end, a 1.6 ms hard refractory period was applied over the events, and the position of the maximum value was recorded as the event time.

### Detection of the position of units using correlated activity at the neighboring channels

One can expect to detect a correlation between the activity of two channels during spike events if the spiking unit is located between these channels. Also, if the unit is not located between these channels *e.g*., at the far end of one of these channels, the activity should not be correlated as the distant channel should be measuring the noise during event times and the noise should average itself out. These arguments suggest that we can approximate the position of the units using the correlation between the channels. Accordingly, peak amplitudes of the events were collected from the channel that the unit belongs to and all the surrounding channels which constitute a 3 x 3 grid together. Then the position of the unit is approximated by averaging the channel positions weighted with the absolute value of the mean of the peak amplitudes

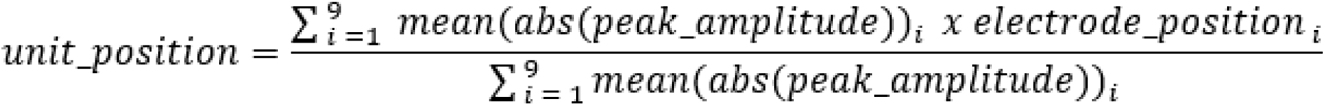

Since some of the channels can capture the waveform in a reversed form, summing up peak amplitudes may sum up to negative values or even 0. Absolute value prevents unphysical results sourcing from small or negative normalization constant at the denominator.

### Elimination of units with non-spikelike waveform

The validity of the resulting units was evaluated according to three conditions. First, units with negligible activity, less than 100 spikes during the recording, were discarded. Afterward, units with a symmetric mean waveform were eliminated. This is the same argument we used during pre-processing. Units were assumed to be symmetric if the correlation between the halves of the waveform was greater than 0.99, and the absolute value of the difference between the halves was less than 2.2e-5. This time no normalization was applied before using the symmetry argument and the center 2 ms part of the waveforms was focused during the elimination. Lastly, units with large noise were also discarded due to their unreliability. We set a lower and upper threshold for the noise and narrowed down the region we inspect to the center 1.2 ms. Any unit with the sum of its standard deviation greater than 2 times or less than 0.1 times of the sum of its mean over its waveform was eliminated. In the end, all results were saved under the channel to which the unit was assigned to.

### Analysis

All the data were labeled with which week they are recorded and with how much cell density the culture initiated with. To evaluate the significance for the findings, Wilcoxon rank sum test applied throughout the analysis. We used the ‘ranksum’ function of MATLAB for the rank sum test.

### Firing rate

Firing rates were found by dividing the number of events by the recording length for each unit. The distribution of firing rate is generated with logarithmic bins = 10^(−2:0.2:1.4).

### Pairwise correlation

To calculate the correlation coefficients between the units we generated binarized time series with various resolution values ranging from 2 ms to 50 ms. The percentage of the events represented for each resolution value is given in supplemental Figure 3. We chose an 11ms resolution which represents more than 95 percent of the events. As a binary value, 1 was assigned to a time point if there is at least one spike within its range, and 0 otherwise. Time series are convolved with a normal distribution, ranging from −10 to 10, centered at the origin and with a standard deviation equal to 10. ‘conv’ and ‘normpdf’ functions of MATLAB were used during the procedure. To calculate the correlation coefficient, the ‘corrcoef’ function of MATLAB was used. To form the correlation distribution, we collected the coefficients from the upper triangle of the correlation matrix. We generated the histogram of these coefficients using the ‘histcounts’ function of MATLAB with 0.05 bin size between −1 and 1.

### Count and word entropies

To quantitatively evaluate the distribution of the network events we facilitated entropy measures. During this analysis, we used binarized time series with an 11ms time resolution that ensured that 95% of bins had only 1 spike. In the remaining 5% of bins that contained more than one spike, we represented this as 1 (irrespective of the number of events). We first compared two variations of entropy namely count entropy and word entropy. Count entropy works on the distribution of the number of simultaneous events. Differently, word entropy works on the distribution of the patterns occurring within the network. Count entropy cannot differentiate the source of the event, but word entropy can. Since we were working with binary data, we used the definition of the Shannon entropy (the information-theoretic entropy):

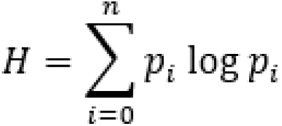

where H is the entropy and p_i_ is the probability of observing the count_i_ for count entropy and pattern_i_ for word entropy, and the sum runs over all possible counts and patterns.

Since word entropy carries the information about the source, oftentimes it becomes impractical to calculate the word entropy directly. Also, one cannot compare the entropy of the recordings with different population sizes. To solve these issues, we subsampled the population of units and calculated the entropy values over these subsamples. The size of the subsampling changed for values k = 2, 4, 6, 8, 10, 12, 14 and analysis was found to be unaffected by the sampling size. However, since increasing the subsampling size requires us to drop the recording with a population size less than the subsampling size, we kept k = 6 for the rest of the analysis to have a more generalizable result. The number of resampling is determined by the size of the population of the units recorded. If population size n has more than 50 combinations with size k (n choose k > 50) subsampling was repeated 50 times. Otherwise, subsampling is repeated n choose k times. We generated the histogram of the entropy values using the ‘histogram’ function of MATLAB with logarithmic bins from 10^-2 to 10 with 0.1 increase in the exponential as step size.

### Maximum entropy

We used Ori Maoz’s maxent_toolbox for the maximum entropy calculations (Maoz and Schneidman 2017). This package generates the maximum entropy distribution for a given training data utilizing the set of constraints applied. The generated distribution has the minimal amount of structure other than the constraints we applied. It is useful for detecting how much of the structure of the data is sourced from the constraints we chose.

We subsampled the units from recording with k = 6. The number of resampling was determined the same way with the former section. We used firing rate, and pairwise interactions parameters as a constraint in our analysis. The max_ent package first calculates the statistical features, mean firing rate <sigma_i> and mean pairwise interaction <sigma_i * sigma_j> and infers the parameters, and maximizes the entropy with the following expression.

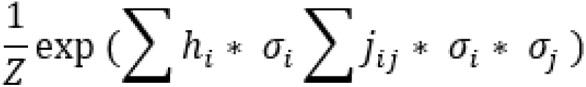

The package generates synthetic data using the inferred parameters and compares it with the empirical data. For this purpose, at the beginning, the algorithm splits the data into two as train and test sets. Results were evaluated by comparing the empirical and the synthetic data using Kullback-Leibler divergence (KL-D). KL-D measures how dissimilar the two distributions are using the following formula.

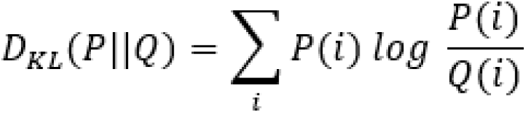

Where P and Q are the word probabilities for synthetic and empirical data respectively (Kullback and Leibler 1951).

Distribution of KL-D values are generated with the ‘histogram’ function of MATLAB with logarithmic bins from 10^-5 to 10 with 0.2 step size on the exponent of 10. The distribution of the parameters were generated the same way, with bins 0:0.5:10 and −10:0.75:5 for h and j terms, respectively.

### Constructing the functional connectivity graph

To visualize and understand the connectivity structure of the network we utilized the firing rate and pairwise correlations. Connection strengths between the units are derived from the pairwise correlations. The connection strength inferred from correlations was evaluated as an undirected relationship. Correlation values around zero are assumed to imply independence between node pairs and neglected. Correlation values greater than 0.2 assumed to imply connection between node pairs and the increase in correlation was directly taken as the connection strength between the node pairs. Graphs representing the network structure visualized using these arguments. Node sizes are determined by their firing rate. They increase with the increasing firing rate. Also thickness of the connection between node pairs is a function of correlation, and it increases with increasing correlation values. The position of the nodes is found as described in ‘the Spike Sorting Using Spatial Information’ section.

To get the distances between the connected units, Euclidean distance was used. The distribution of the physical distances of the functionally connected units was generated with the ‘histogram’ function of MATLAB with logarithmic bins from 10^-4 to 10^0.8 with 0.2 step size in the exponent. Cumulative sum of distances generated using this histogram. The following heat maps in Figure 7 C1-2 were generated with logarithmic bins. We used the ‘linspace’ function of MATLAB for generating distance bins from log10(1e-4) to log10(4) in 30 steps and correlation bins from log10(1e-1) to log10(1) in 30 steps. Resulting color scale is also represented in the log scale.

## AUTHOR CONTRIBUTIONS

R.S., M.C.M., and K.P. conceptualized the study and the experiments, R.S. performed the experiments and curated the data, Y.S.U., and K.P. conducted the formal analysis and generated figures for the manuscript. Y.S.U., and K.P. wrote the original draft, R.S., and M.C.M. reviewed and edited the manuscript.

## ACKNOWLEDGMENTS

We are grateful for the funding provided to M.C.M. by the Larry L. Hillblom Foundation and the donation from Agilebio to R.S.. K.P. was supported in part by NIH R01MH113924, R01NS135763, P50HD103536, NSF 1749772, the Cystinosis Research Foundation, a University Research Award for the University of Rochester, and a Del Monte Pilot Grant. R.S. is a CNRS researcher.

## SUPPLEMENTAL INFORMATION

**Supp. Figure 1.**
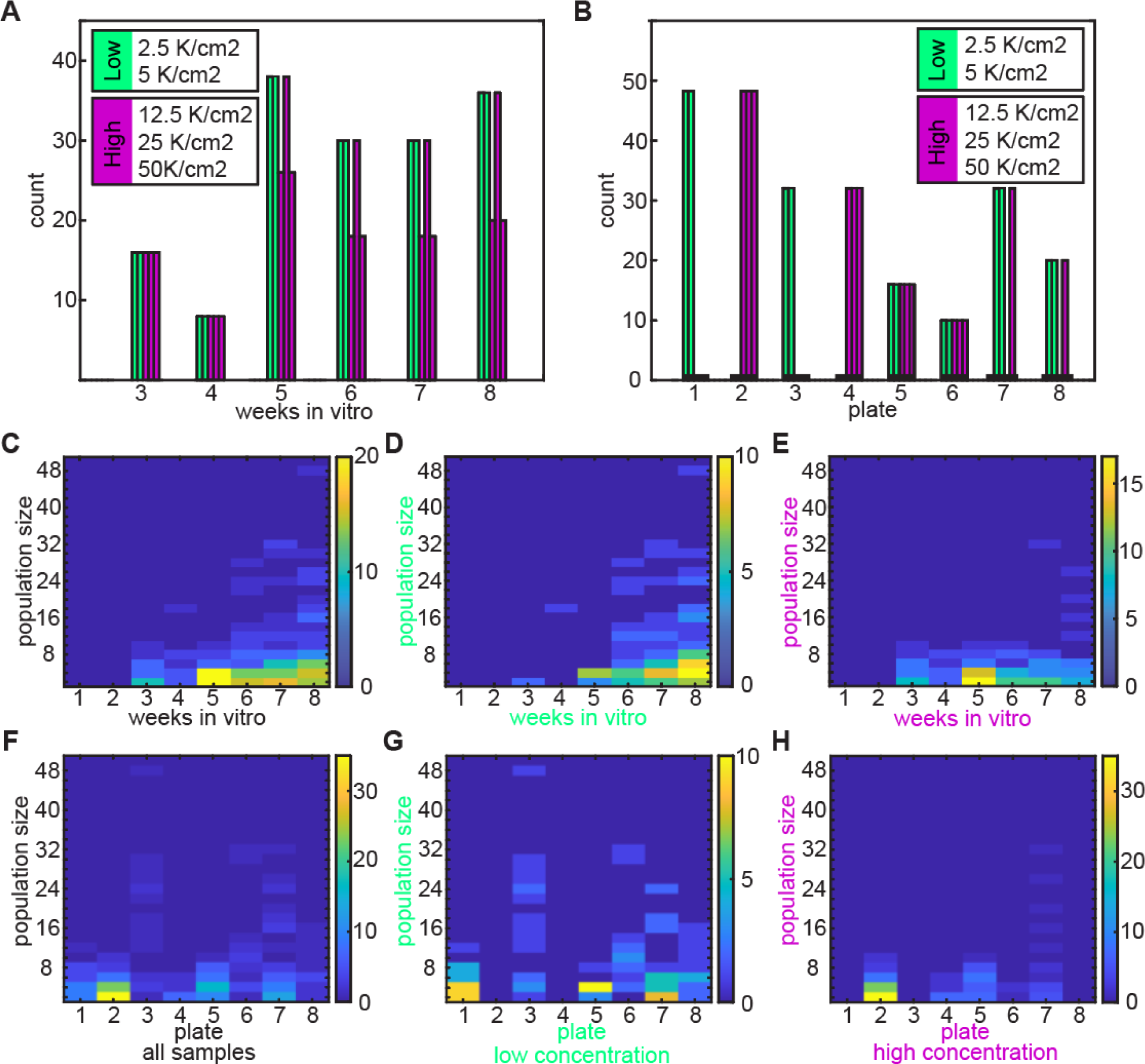
Description of recording weeks and plates. (A) Number of recordings run each week for low- and high-density cultures. (B) Number of recordings made with each plate. (C) Number of recordings with given population size and weeks *in vitro*. (D) Number of recordings with given population size and weeks *in vitro* for low-density cultures. (E) Number of recordings with given population size and weeks *in vitro* for high-density cultures. (F) Number of recordings with given population size and plate. (G) Number of recordings with given population size and plate for high-density cultures. (H) Number of recordings with given population size and plate for high-density cultures.

**Supp. Figure 2.**
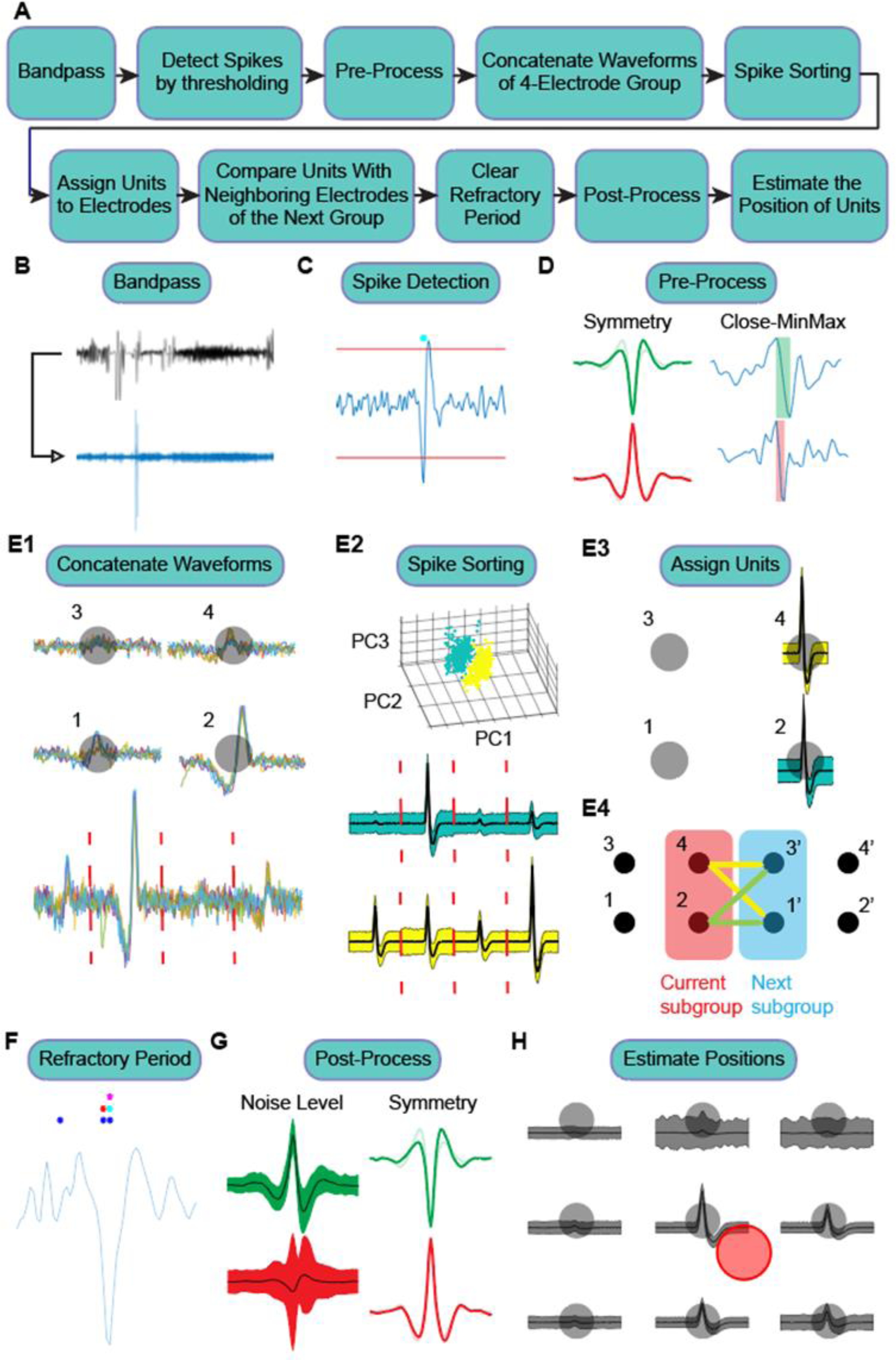
Spike sorting procedure. (A) Overview of the detection procedure. (B) Example raw and bandpassed recording. (C) Spikes were detected with thresholding. (D) Detected spikes eliminated according to two criteria before they were passed to the sorting procedure. Symmetric waveforms were eliminated due to their non-biological form. Also, waveforms with close peak to trough distance were eliminated since sudden jumps in the raw recording turns into a waveform like shape with close min-max (E1) Sorting procedure was run over groups of four channels. Their waveforms were concatenated for all detected spikes. (E2) New waveforms were categorized according to their low dimensional representation as defined with principal component analysis (PCA). (E3) Detected units assigned to the channel where it has the maximum amplitude. (E4) We checked whether the detected units have higher amplitude in a neighboring channel of another group. If that was the case, the unit was removed from the current group and sorted with the next group. (F) Within a group multiple channels can detect the same spiking activity. When we were concatenating the waveforms, we gathered all detected spike times together. Due to noise, the same spike was detected at different locations by different channels. These waveforms were compared with each other using Pearson correlation. If the correlation value between them was 1, we eliminated either one of them. We made sure that there was no detected spike within the refractory period of each other. (G) Detected units were eliminated if either their noise level was too high or they had a symmetric waveform. (H) Physical location of the detected units was determined by weighting the channel locations with the amplitude of the unit waveform in that channel.

**Supp. Figure 3.**
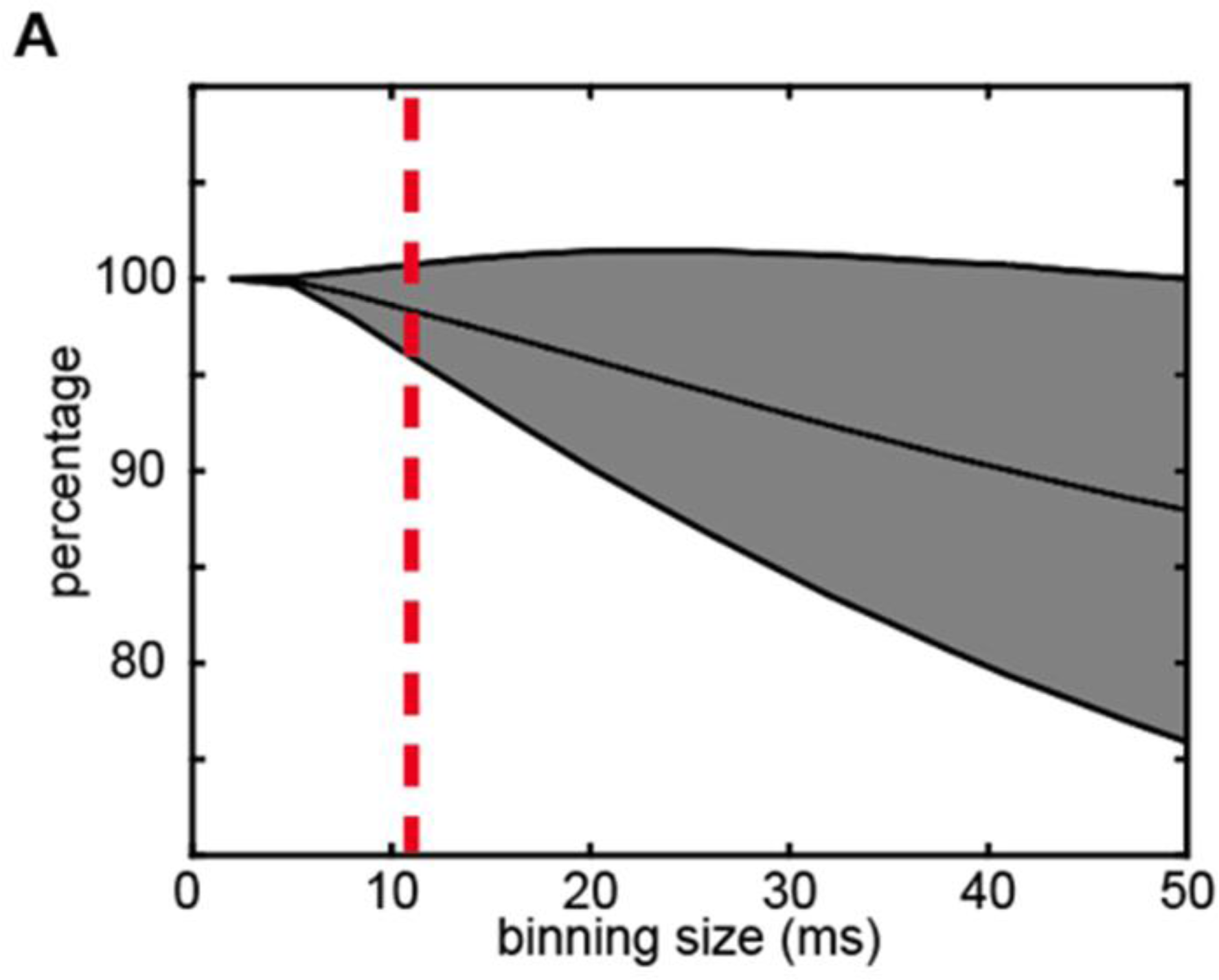
Binning size versus percentage of spikes represented. (A) Percentage of the spikes represented changes for each recording. Gray area is showing the region where the mean ± standard deviation of spikes represented, for all recordings. Increasing binning size decreases the percentage of spikes represented. Throughout the analysis we applied 11 ms binning which accounts for more than 95% of the spikes for all recordings.

